# Capacity of oil palm (*Elaeis guineensis* Jacquin) male inflorescences to host *Elaeidobius kamerunicus* Faust (Coleoptera: Curculionidae: Derelominae). Comparison between two cultivars in northern Sumatra

**DOI:** 10.64898/2026.10.12.758427

**Authors:** L. Beaudoin-Ollivier, E. Freyssinet, R. Rahman Moeis, H. Rey, D. Afandi, I. Syahputra, C. Madec, Y. Dumont

**Affiliations:** CIRAD, UMR ABSys, F-34398 Montpellier, France; ABSys, Univ Montpellier, INRAe, SupAgro Montpellier, France; Université Jean Monnet, Saint-Etienne, France; Bogor Agricultural University, Bogor, Indonesia; CIRAD, UMR AMAP, F-34398 Montpellier, France; AMAP, Univ Montpellier, CIRAD, CNRS, INRAE, IRD, Montpellier, France; PT Socfindo SSPL, Bangun Bandar, 20991 Sumatera Utara, Indonesia; Palmelit S.A., boulevard de la Lironde, 34398 Monferrier-sur-Lez; CIRAD, UMR AMAP, Pole de Protection des Plantes, F-97410 Saint-Pierre, France; Department of Mathematics and Applied Mathematics, University of Pretoria, 001 Pretoria, South Africa

**Keywords:** *Elaeidobius kamerunicus*, male inflorescence, anthesis, inflorescence carrying capacity, pollination, super-male palm

## Abstract

During the 1980’s, *Elaeidobius kamerunicus*, one of the oil palm pollinator species in Africa, was successfully introduced in Indonesia from Cameroon to improve oil palm production and reduce assisted pollination. It is now well known that *E. kamerunicus* completes its entire life cycle on the anthesis of the male inflorescences. However, for a long time, oil palm breeding programs have favoured the development of female inflorescences for production purposes, to the detriment of male inflorescences. The objective of our study was to explore the interactions between the weevil population, at immature stages, and male inflorescences, produced by two different cultivars (a super-male (SM) cultivar and a highly female and productive experimental cultivar (EXP) of five years old) during anthesis and post-anthesis stages. Morphological characteristics of spikelets of inflorescences were also studied and compared. We analysed field and laboratory data, collected in North Sumatra (Indonesia) from April to June 2019. The mean number of spikelets is lower in EXP palms (126±16) than in SM palms (213±15). There are more abnormal spikelets in SM palms (126±77 and 100±73 for SM-SD and SM-DD cultivars, respectively), than in EXP palms (33±23). However, all these differences do not affect either male inflorescences number of flowers or their carrying capacity that contributes to *E. kamerunicus* development. The number of male flowers per inflorescences is almost the same for both materials, and about 10% of flowers were occupied by *E. kamerunicus* immature stages. Despite noticeable morphological differences, no major differences were found between male inflorescences of EXP and SM cultivars. We will discuss our results from pollination point of view.

## Introduction

The oil palm *Elaeis guineensis* Jacquin 1763 is a tropical plant of economic interest and the world’s leading source of vegetable oil. Fruit set rate and then production of well-formed fruits in a bunch depends mainly on pollen transported by *E. kamerunicus (EK)* introduced in Southeast Asia in the 1980’s.

The oil palm is a monoecious perennial herb and bears asynchronous male and female flowers on separate inflorescences. *E. kamerunicus* are involved in a specialized mutualistic relationship with their host plant (Syed, 1979). The male inflorescence in which the insects develop is highly attractive to *E. kamerunicus* and, through an olfactory deception phenomenon, they visit the female inflorescence on which they deposit pollen. *E. kamerunicus* was found to be highly specific to the genus *Elaeis* (Syed, 1979; Adaigbe et al., 2011). Visitors to oil palm inflorescences are attracted by rewards such as food, shelter and oviposition sites.

The male inflorescences are composed of digitiform spikelets carrying between 1,400 and 1,500 flowers (Hartley, 1988; Turner and Gillbanks, 1974). According to (Jacquemard, 2011), there are between 100 and 300 spikelets per inflorescence. Spikelets measure from 12 to 20 cm in length (Hartley, 1988; Prasetyo and Susanto, 2012). Anthesis progresses acropetally in male inflorescences (Tandon et al., 2001) over a period of 4-5 days (Dhileepan, 1994). During the anthesis stage, the flowers progressively open from the bottom to the top of the inflorescence until all florets in spikelets are opened (Chiu et al., 1986; Li et al., 2019) emitting a strong aniseed fragrance. The anthesis stage lasts about five days (Corley and Tinker, 2016; Forero et al., 2012), followed by the post-anthesis stage, marked by flower withering (Chiu et al., 1986).

*E. kamerunicus* is holometabolous, undergoing complete metamorphosis inside the staminate flowers (Siswanto and Soetopo, 2020). The adult chew anther filaments or anther tubes of opened male flowers (Syed, 1982). They mate at any time of the day, usually 2 to 3 days after emergence. However, some individuals mate sooner (Syed, 1982). The female lays eggs inside individual male flowers of the anthesis male inflorescence close to the superior part of the anther (Syed, 1982). Eggs are whitish, oval with a smooth, delicate chorion, 0.65 mm long and 0.4 mm width (Syed, 1982). *E. kamerunicus* laid, on average, 57.64 eggs in male flowers (Wahyuni et al., 2020). When hatching 1-2 days afterwards, neonate larvae feed on flowers. The larva is translucent and grey and moult two times before to reach the prepupae stage (Syed, 1982). Often the egg and first two larval stages of *E. kamerunicus* are completed within 2-3 days (Syed, 1982). There are 3 larval instars. During the last larva instar, it perforates the perianth solid base to move to the neighbouring flower on which the larva feeds. One larva is able to feed on up to six male flowers to perform its development (Syed et al., 1982). The prepupae stop feeding and moving to become opaque and yellow in colour. Pupae are immobile within the flower, yellow in colour and have morphological traits of the future adult. The life cycle lasts about 10 days in the male inflorescence (Syed, 1982; Tuo et al., 2011). However, among all works done on *EK* and male inflorescences, there is no information about the male inflorescence capacity to “produce” emerging weevils. In other words, we found that knowing the average carrying capacity of male inflorescences, for a given material, is necessary to estimate the average density of weevils per ha.

*E. kamerunicus* was never observed on male spikelets before anthesis but arrived in large numbers as soon as a few florets had opened at the base (Syed, 1982). The greater number of insects congregate at male inflorescence corresponding to the greater number of volatiles compounds release. (Mohamad et al., 2024) reported 4,267 *E. kamerunicus* captured on sticky traps placed above male inflorescences in anthesis.

The current concern in the oil palm sector is regarding the decrease in fruit set that may be affected by *E. kamerunicus* population density (Prasetyo et al., 2014; Zulkefli et al., 2020). From (Kouakou et al., 2018), the fruit set is positively correlated to the *E. kamerunicus* population visiting the male and female inflorescences. Thus, the weevil population plays a crucial role in the pollination of oil palm.

In Indonesia, the *E. kamerunicus* population is fluctuating due to several factors such as unfavourable climatic conditions or a low density of male inflorescences (Syed and Saleh, 1988; Li et al., 2019). Environmental stresses (climatic, hydric…) usually trigger the production of male inflorescences with a lag of 14 months (Turner and Gillbanks, 1974). However, climatic conditions in Indonesia are often favourable and oil palm produce more female inflorescences than male inflorescences. Naturally, the sex determination of oil palms, which is described as the ratio of female to overall inflorescences produced by a palm, is most affected by climatic variables in which drought increases the ratio of male inflorescences (Yousefi et al., 2020). Additionally, oil palm breeding programs have favoured high-yielding planting material with a higher ratio of female inflorescences at the expense of male ones (Corley and Tinker, 2016).

However, more female inflorescences result in less available pollen and fewer weevil breeding sites, potentially reducing pollination (Rao and Law, 1998; Bulgarellil et al., 2002). As a consequence, the natural pollination can be poorer in improved oil palm cultivars (Syed et al., 1982; Nurul Fatihah et al., 2019). For this reason, « super-male » palms have been developed and tested to increase the number of male inflorescences and to maintain *E. kamerunicus* population above the minimum threshold needed to effectively pollinate female inflorescences.

The strategy of integrating planting material with a large number of male flower spikelets may have a positive effect on the population of *E. kamerunicus* (Teo, 2015; Wahyuni et al., 2020).

We present a study done in Indonesia, comparing the potential of male inflorescences to host *E. kamerunicus* immature stages for two types of planting material i.e., a highly female and productive experimental material and a super-male material. Observations were carried out at different scales, at the spikelet and inflorescence level. The objectives of the study were (1) to compare morphological characteristics of male inflorescences for the two different planting materials (2) to estimate and compare the weevil’s carrying capacity of each material to further determine whether they influence the population abundance of immature *E. kamerunicus*.

## Material and methods

### Field site

The present study was carried out at Bangun Bandar estate (3°17’°03.12 ‘’N 99°01’57.73’’E) at SSPL, Socfindo Seeds Production Laboratory, located at about 80 km from Medan on Sumatra Island in Indonesia. North Sumatra region is characterized by a dry season from May to September. However, the difference with the rainy season is not very pronounced, no water deficit is usually observed. Field and laboratory observations were made between April 1^st^ and June 20^th^, 2019, during a relatively dry season. Rainfall data recordings did not exceed 400 mm during that period.

The observation site was a standard 64 ha plantation located on tropical mineral soils, surrounded by rubber and young oil palm plantations to limit undesired effects of neighbouring pollen.

### Planting material

Observations were carried out on three oil palms at 5-years after planting, every week, from April 6^th^ to June 14^th^, 2019, during the “dry season”. Two different planting materials were compared: (i) a super-male cultivar consisting of the two clones SM-1 and SM-2 which were developed by Palmelit SAS company (France) to produce a constitutively high number of male inflorescences (Dumont et al., 2026), and (ii) an experimental cultivar (EXP) corresponding to a highly female and productive planting material (Cochard et al., 2005).

EXP palms were planted at a standard density of 143 palms / ha (single density (SD)), whereas super-male palms (SM-1 and SM-2) were planted at 3 different densities:

(1) 143 palms / ha with 138 EXP and 5 (permanent) SM, in one row out of six, every five palms. The 5 “permanent” SM palms were planted in the place of EXP palms (single density (SD)).
(2) Following the same planting design (as the one described in 1) but with 5 additional SM (within the windrow, in double density (DD)) planted in one windrow out of 6 and three rows after the one with the permanent SM.
(3) Following the same planting design (as the one described in 1) but with 10 additional SM (within the windrow, double density (DD)) planted in one windrow out of 6 and three rows after the one with the permanent SM.

For each individual series of observation, three inflorescences were randomly selected in the field, five days after the beginning of the anthesis, after *E. kamerunicus* females laid their last eggs: one EXP inflorescence and two SM inflorescences, planted in single and double density respectively. Each series lasted six successive days, from the post-anthesis stage and before to first imago emergences. A total of nine series of observations were made, and related data were recorded from April to June 2019.

Spathes around inflorescences were removed to access spikelets daily and collect them at any level on the inflorescence. Four spikelets were collected daily for five days (before the beginning of the first imago emergence) between 07:00 a.m. and 08:30 a.m., by cutting them at 3 different levels: bottom (BOT), center (CEN) and top (TOP) of the inflorescence using a shear. Two of the spikelets were collected in the front site and two others at the backside of the inflorescence. For each level, spikelets were numbered 1 and 2 (front collection), 3 and 4 (backside) and surrounded by three different colour scratch tape representing the level on the inflorescence. The spikelets were stored in a plastic bag for a short transfer from the field to the laboratory. Planting material, density of the SM, exact location in planting block (row and palm numbers), were mentioned on the plastic bag. The dissection of spikelets occurred the same day than collection to avoid them to dry off. For each series, a total of 60=5*3*4 spikelets were studied.

The total length for each normal spikelet was measured using a metallic roller from the first flowers at the base of the peduncle to the base of the spine of the spikelet (). For abnormal spikelets, i.e., constrictions without flowers, only lengths with flowers were taken into consideration (Figure 2). These refer to utility lengths (Li). The analysis used a generalized mixed linear model following a quasi-binomial distribution to compare the percentage of normal and abnormal spikelets among planting materials.

**Figure 1.**
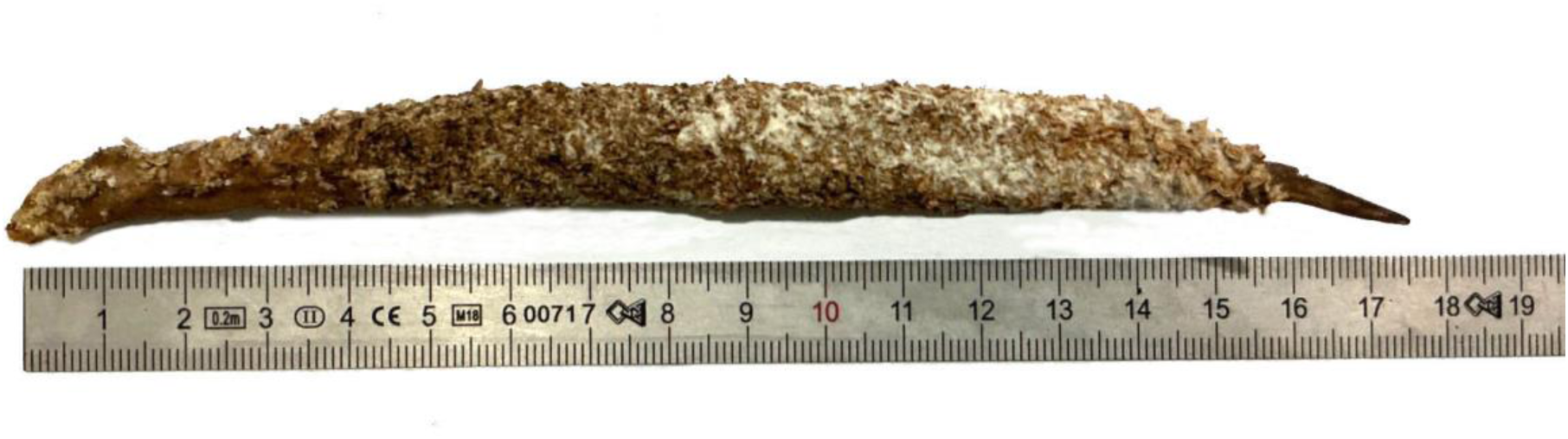
One single male spikelet.

**Figure 2.**
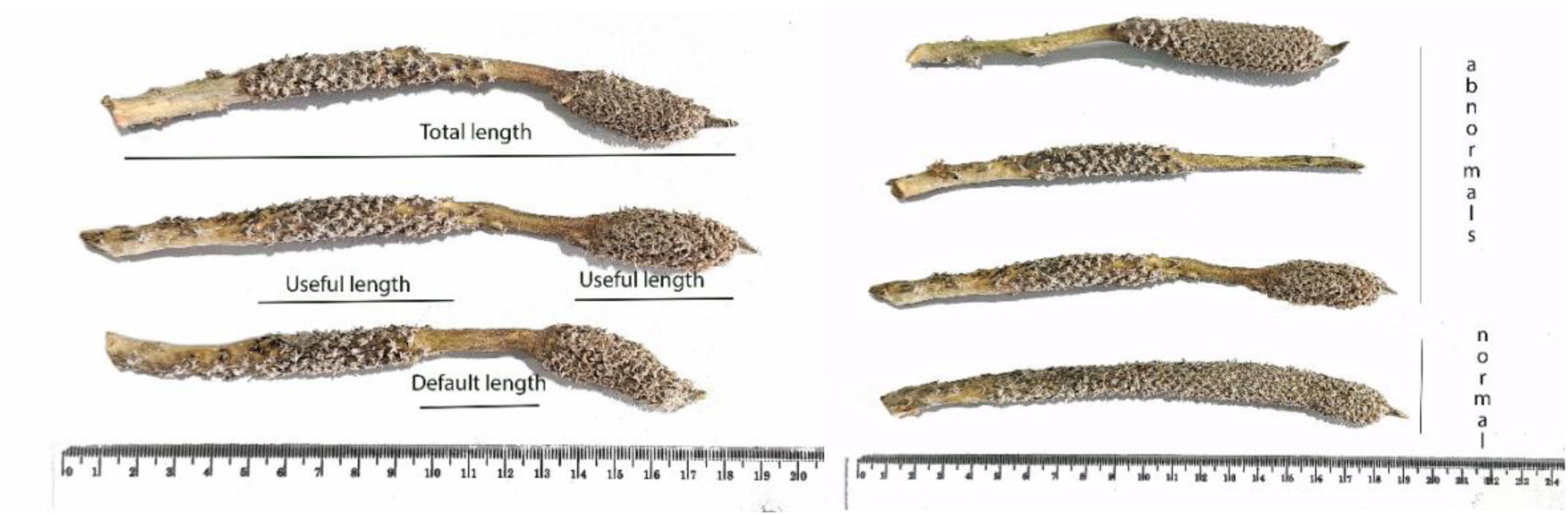
Abnormal spikelet measurement.

Each normal spikelet was cut in three parts (P). For each part, we considered two samples (Sam) (by default 3 cm or less if necessary), one radial (R) and one longitudinal sample (L).

One to three samples measured from 1 to 3 cm (L) for each spikelet were radially cut in three parts (L_1_, L_2_ and L_3_), depending on the spikelet total length and aspect (normal or abnormal) (Figure 3). Each of the three samples were longitudinally cut in four strips named L_1a_, L_1b_, L_1c_ and L_1d_ (Figure 4). Each abnormal spikelet was considered depending of its individual morphology (Figure 2). The number of flowers is counted for each strip (NF_St_). The different immature stages of *E. kamerunicus* (eggs, larva, prepupae and pupae) occupying individual flowers were counted for each strip. Three additional radial cuts (of 1cm long) allowed to assess the theoretical number of flowers per spikelet by using the average measurement of their two diameters (NRF) (Figure 3 and Figure 4). The observations were done under a stereomicroscope (Bresser, Science ETD-201). Similar approach was used by (Prasetyo et al., 2013) to calculate the number of *E. kamerunicus* immature stages from samples of male inflorescences spikelets.

**Figure 3.**
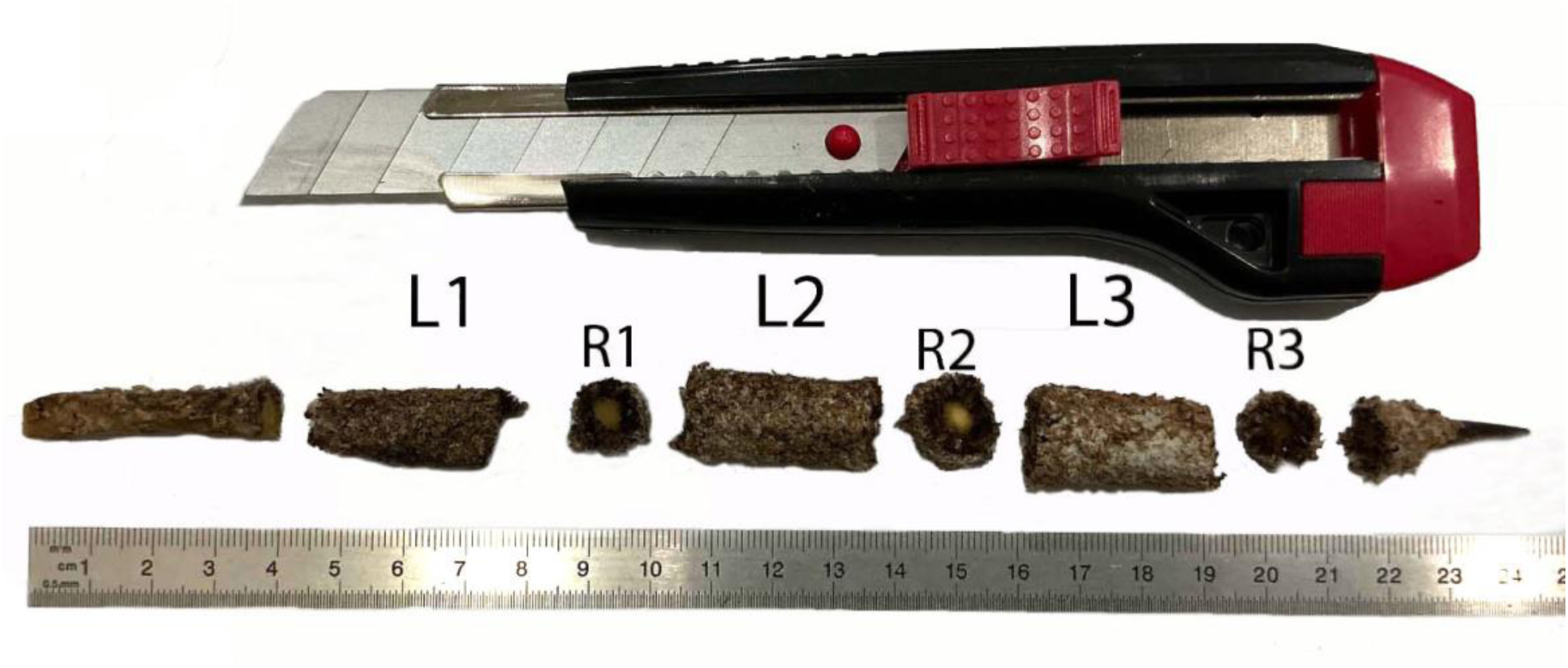
Radial samples along a spikelet (3 cm and 1cm respectively).

**Figure 4.**
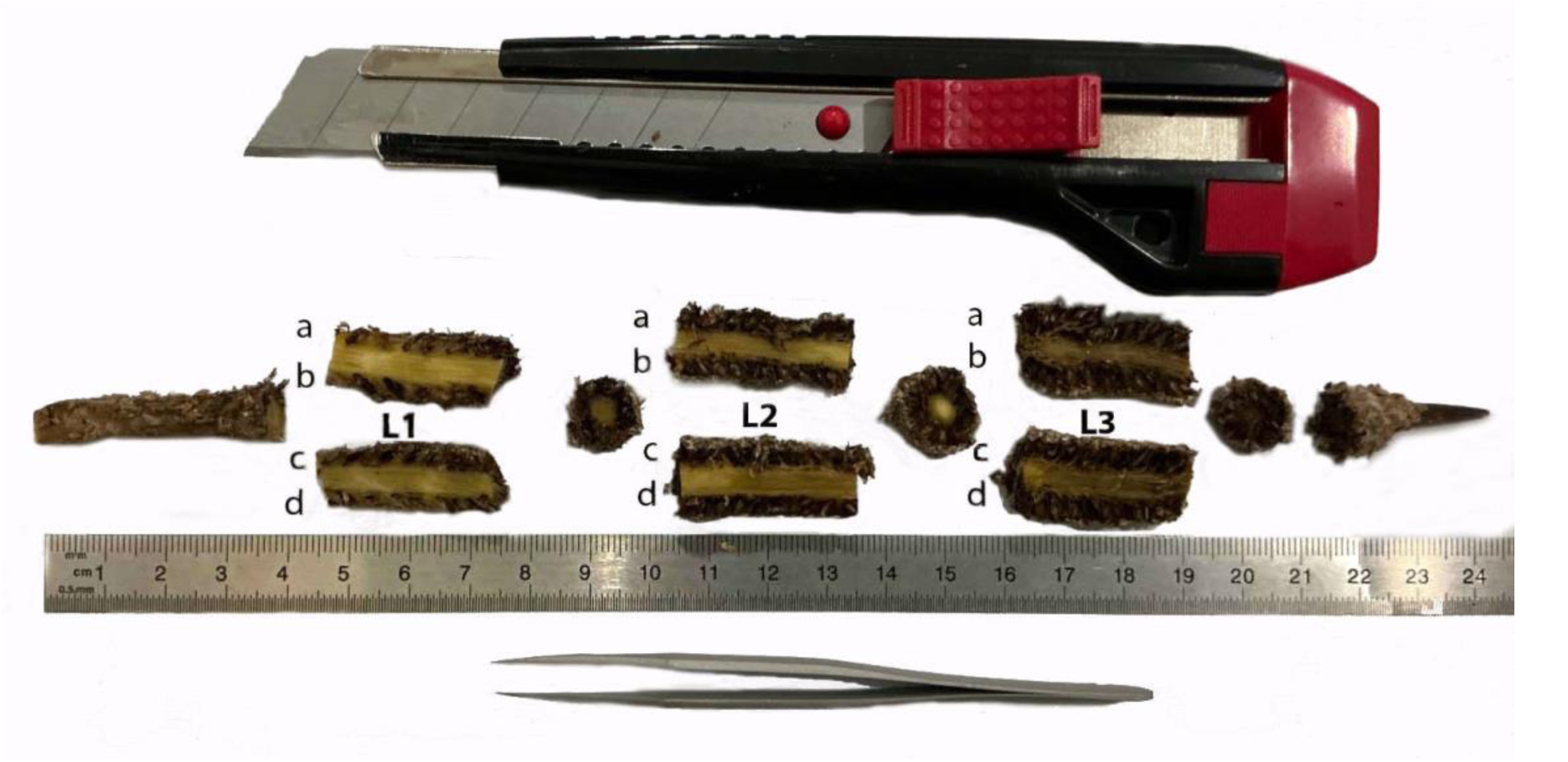
Longitudinal and radial samples along a spikelet.

After five days and the last collection of spikelets, all remaining spikelets were cut, counted and measured per level (bottom, center and top) for each of the three inflorescence types, EXP, SM-SD (super-male in single density) and SM-DD (super-male in double density).

A total of 9 series were studied including 9 inflorescences from EXP palms, 7 from SM-SD (2 were lost due to rat attacks), and 9 from SM-DD. Around 60 spikelets were dissected per series from each inflorescence while all spikelets, from all studied inflorescences, were measured (Table 1).

**Table 1.** Summary of collected data.

| Series | EXP | SM-SD | SM-DD |
| --- | --- | --- | --- |
| Nb of inflorescences | 9 | 7 | 9 |
| Nb of measured spikelets | 1,152 | 1,504 | 1,903 |
| Nb of dissected spikelets | 486 | 378 | 492 |

Based on measurement described above, for each spikelet, the average number of flowers per cm (*NF_st_*) was estimated as the average number of flowers (on all 4 strips) divided by the average length of strips, *L_St_*,

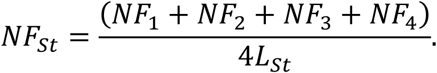

Then, all available flowers on one inflorescence were estimated using the following formula,

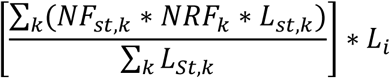

Where k is the number of spikelets studied on each inflorescence, and

- NF*_st_* is the average number of flowers related to the longitudinal sample.
- NRF is the average number of radial flowers
- *L_st_*, is the average length of each longitudinal sample
- *L_i_* is the (utility) length of the spikelet.

### Statistical analyses

Statistical analyses were carried out with R software (version 4.4.2, R Core Team 2024). All data are expressed as mean ± 95% asymptotic confidence interval.

Depending on the type of data, we considered different type of statistical models. Linear Models (LM), linear mixed-effects model (LMM), and Generalized Linear Models (GLM) were built. Models fitting were conducted using the function “glm” from the “stat” package for LM and GLM, and “lmer” from the “lme4” package for LMM. Model assumptions, including homoscedasticity, normality, and independence of residuals, were checked graphically. Significance of fixed effects was tested by Deviance tests via the “Anova” function from the package “car”. When effects were significant (*p* < 0.05), pairwise post-hoc tests were performed using the “emmeans” function from the “emmeans” package.

#### Effect of the planting material on the Number of spikelets per inflorescence

To analyze the effects of the planting material on the Average Number of spikelets, we considered a Generalized Linear Model (GLM) with a quasi-Poisson distribution. Then, an Anova II test was then used to assess the significance of the fixed effects.

#### Effect of the planting material and the spikelet’s position within the inflorescences on the useful length and the number of (occupied) flowers per spikelet

Estimated useful length, at the spikelet and the inflorescence levels, was analysed using Linear Mixed Models (LMM) including the effects of the material and the position of the spikelet, within the inflorescence, as fixed factors, and the inflorescence as random factor. An Anova III test was then used to assess the significance of the fixed effects, followed by Tukey Post-hoc analysis.

#### Effect of the planting material and the spikelet’s position within the inflorescences on the proportion of Normal and Abnormal spikelets

A Generalized Linear Model (GLM) with a binomial distribution was considered. Then, an Anova II test was then used to assess the significance of the fixed effects, followed by a pairwise post-hoc analysis.

The effect of planting materials on the total useful length, the total default length, and the number of (occupied) flowers per inflorescence was studied using Linear Models and post-hoc analysis.

#### Effect of planting materials on the immature population structure up to 4 days after the end of the anthesis stage

Between the end of anthesis M_5_ (5 days after the start of anthesis) and the following 4 days (M_6_, M_7_, M_8_ and M_9_), all immature stages of *E. kamerunicus* (i.e., eggs, larvae, prepupae and pupae) were estimated using the dissected spikelets, for each planting materials. To compare the pollinator population structures among the three planting materials, Chi-squared (*χ*^2^) tests of independence were conducted separately for each anthesis stage (M_6_ to M_9_), followed by a Pearson standardized residual analysis by anthesis stage, aiming to identify which planting material deviated from the theoretical means for each development stage of the pollinator.

## Results

### Inflorescence’s morphology

According to Figure 5, the number of spikelets was higher for both SM-DD (211 ± 13) and SM-SD (215 ± 17) materials than the EXP material (128 ± 16). The statistical analysis confirmed that the difference between both SM materials and the EXP material was statistically significant (*p* < 0.0001). As expected, there was no statistically significant difference of the average number of spikelets between SM-SD and SM-DD planting materials (*p* = 0.9471).

**Figure 5.**
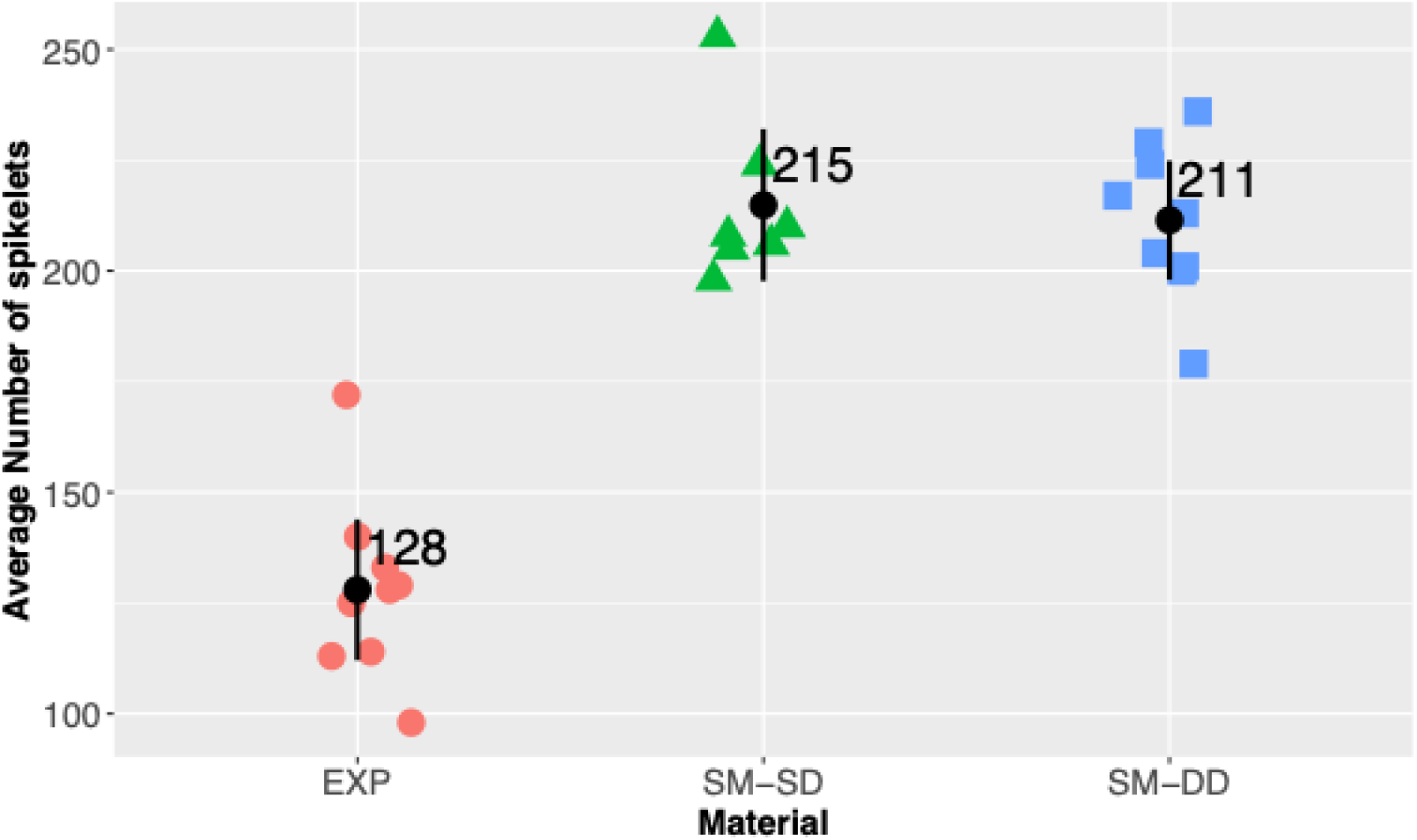
Total number of spikelets (average and confidence interval) for each type of planting material.

In Figure 6, we show the spikelet’s useful length data measured on the 9 EXP, 7 SM-SD, and 9 SM-DD inflorescences, for each position on the inflorescence (bottom, center and top). For all planting materials, on average, spikelets at the bottom are longer than spikelets in the center, themselves longer than spikelets at the top. The EXP material always has the longest useful spikelets length, on average.

**Figure 6:**
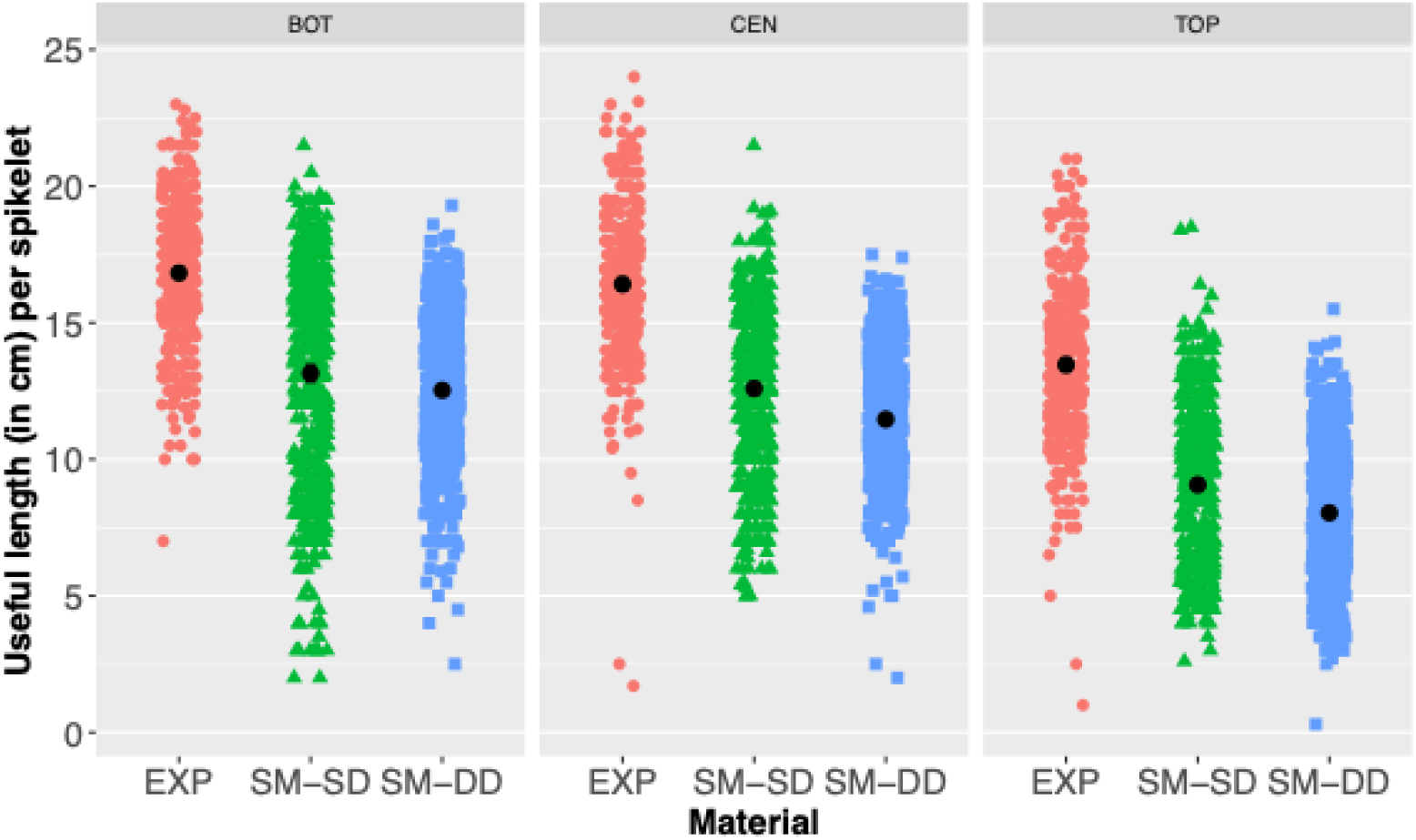
Useful length (in cm) per spikelet measured on the EXP, SM-SD, and SM-DD planting materials, according to their position on the inflorescence, at the bottom (BOT), center (CEN) or the top (TOP).

Spikelet useful length varied significantly among planting materials (*p* = 2.15 × 10^−5^), with spikelets from EXP inflorescences having longer useful length than spikelets from both SM inflorescences (SM-SD and SM-DD). The spikelet position is also highly significant (*p* = 2.2 × 10^−16^). Moreover, a significant interaction between spikelet position and planting material was detected (*p* = 5.10 × 10^−8^). For EXP inflorescences, spikelets from both the bottom and the center had a significant longer useful length than spikelets from the top of inflorescences. However, for SM inflorescences, spikelets from the bottom were significantly longer than spikelet from the center, themselves significantly longer than spikelets from the top.

Based on data presented in Figure 5 and Figure 6, the average total useful length per inflorescence was calculated. Using a Tukey post-hoc test, we showed that there was no significant difference between the EXP and the SM-DD planting materials (Figure 7; *p* = 0.374), and between the SM planting materials (*p* = 0.4913). However, there was a statistical trend between the EXP and the SM-SD planting materials (Figure 7; *p* = 0.058). Additionally, we obtained similar result for the average total default length per inflorescence (Figure 8): no statistical difference between the SM planting materials (*p* = 0.4974) and the EXP and SM-DD planting materials (*p* = 0.3683), and a statistical trend between the EXP and SM-SD materials *p* = 0.0585).

**Figure 7.**
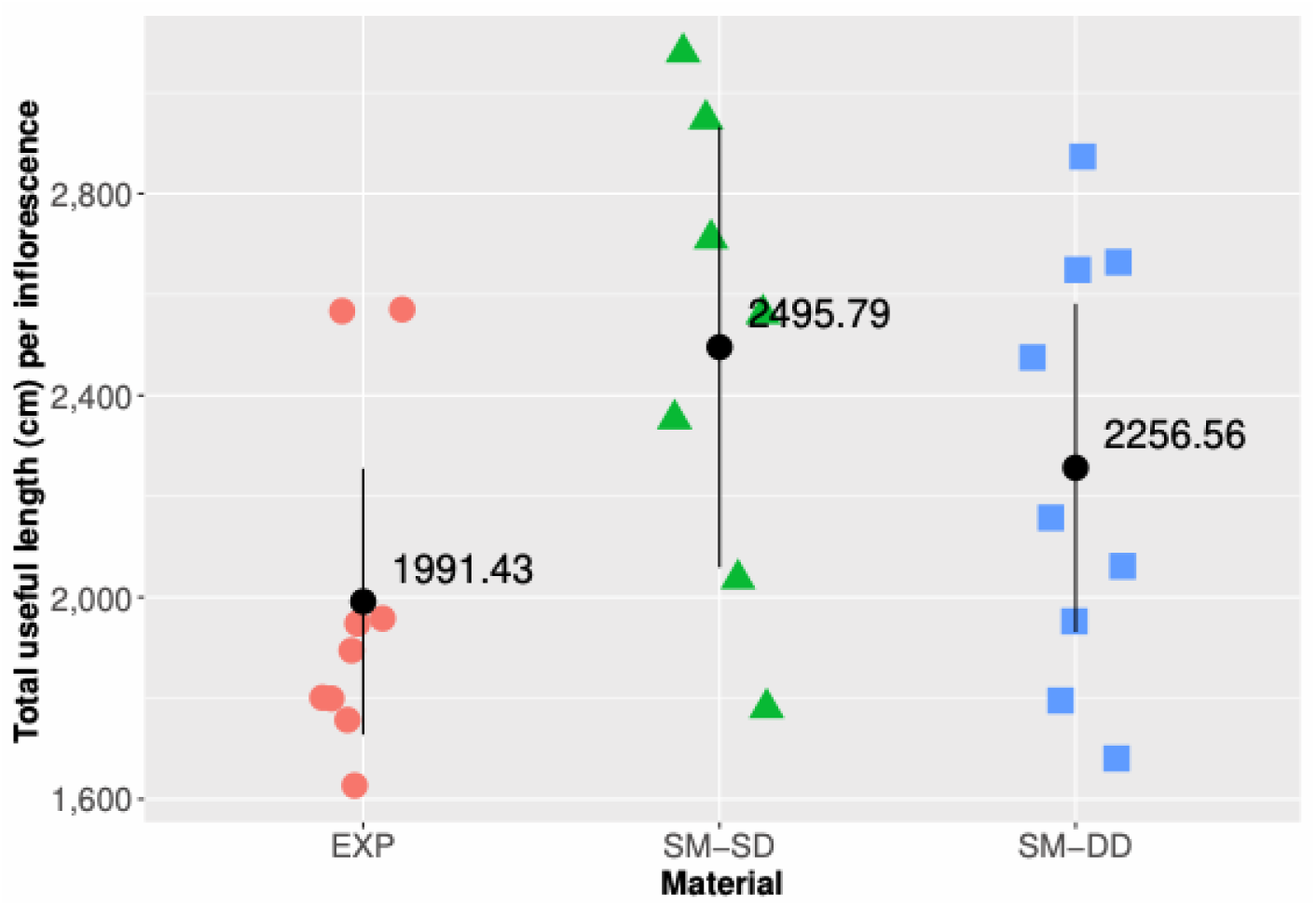
Total useful length (in cm) for each type of planting material.

**Figure 8.**
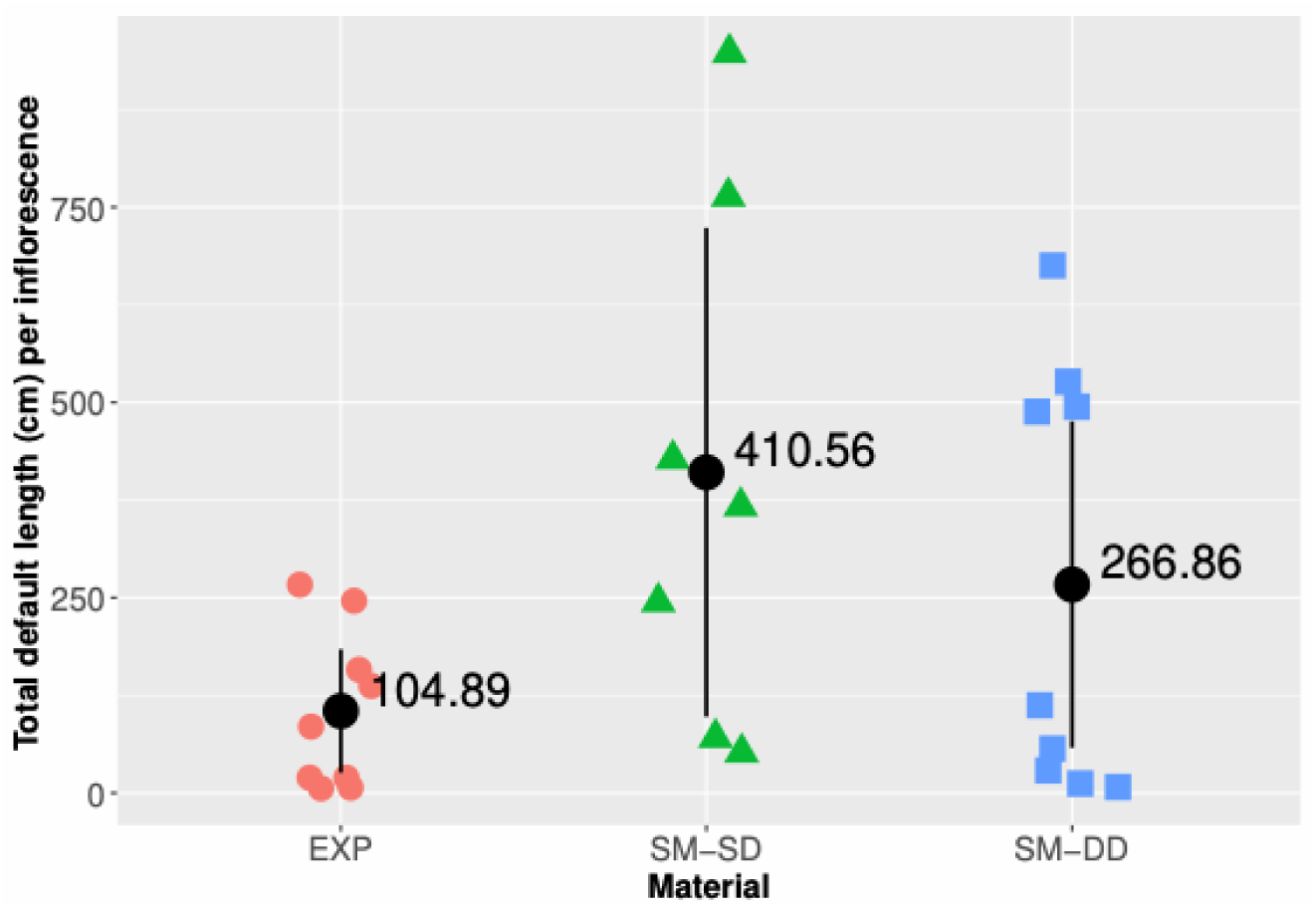
Total length (in cm) of spikelets without flowers.

### Normal and abnormal spikelets

A significant interaction between spikelet position and planting material, on the percentage of normal and abnormal spikelets was detected (Type II Anova, *p* < 0.001). The percentage of abnormal spikelets (Figure 9, in blue) was lower for the EXP material (33 ± 22.62) than for both SM materials (SD, 126 ±76.61, DD, 100 ± 72.54). The proportion of abnormal spikelets was larger at the top of the inflorescence for the SM-SD and SM-DD materials. Then, the pairwise comparison showed that, for each planting material, the position along the inflorescence had a significant impact on the proportion of normal spikelets (*p* <0.001). However, statistically, there was no differences between all material at the center and the top (p>0,29), and no differences between the SM-SD and SM-DD materials whatever the position (p>0,67).

**Figure 9.**
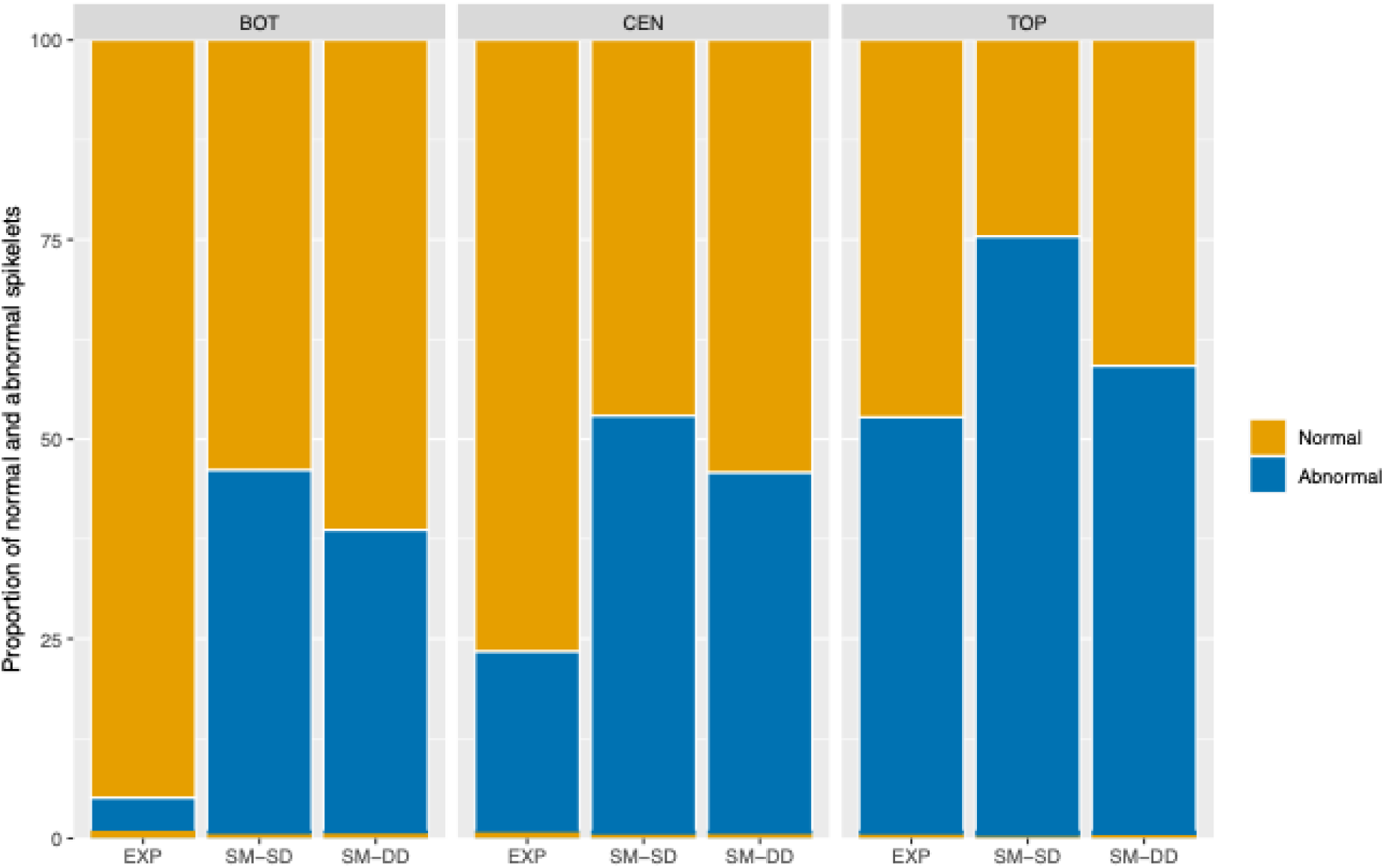
Percentage of normal (yellow colour) and abnormal (blue colour) spikelets depending on the position within inflorescences.

### Number of flowers

The number of flowers per spikelet was significantly affected by the planting materials (*p* < 0.0024), but with no significant differences at the center/bottom of the inflorescence (p>0.21). SM-SD and SM-DD planting materials produced less flowers per spikelets (815 ± 241, and 675 ± 91, respectively) than the EXP planting material (1,349 ±296; Figure 10). A significant interaction between planting material and spikelets position was also found (p < 0.0001). Post hoc comparisons showed no significant difference between both SM materials (*p* > 0.1291), independently of spikelet position. Additionally, no significant difference was found between spikelets from the bottom and the center of inflorescences for EXP and SM materials (*p* > 0.2114), but number of flowers per spikelets differed significantly between the bottom/center and the top of inflorescences for EXP and SM materials (*p* < 0.0001). At the top, they were 20% less male flowers for the EXP material, 50% for the SM-SD material, and 75% for the SM-DD material than at the bottom/center of the inflorescence.

**Figure 10.**
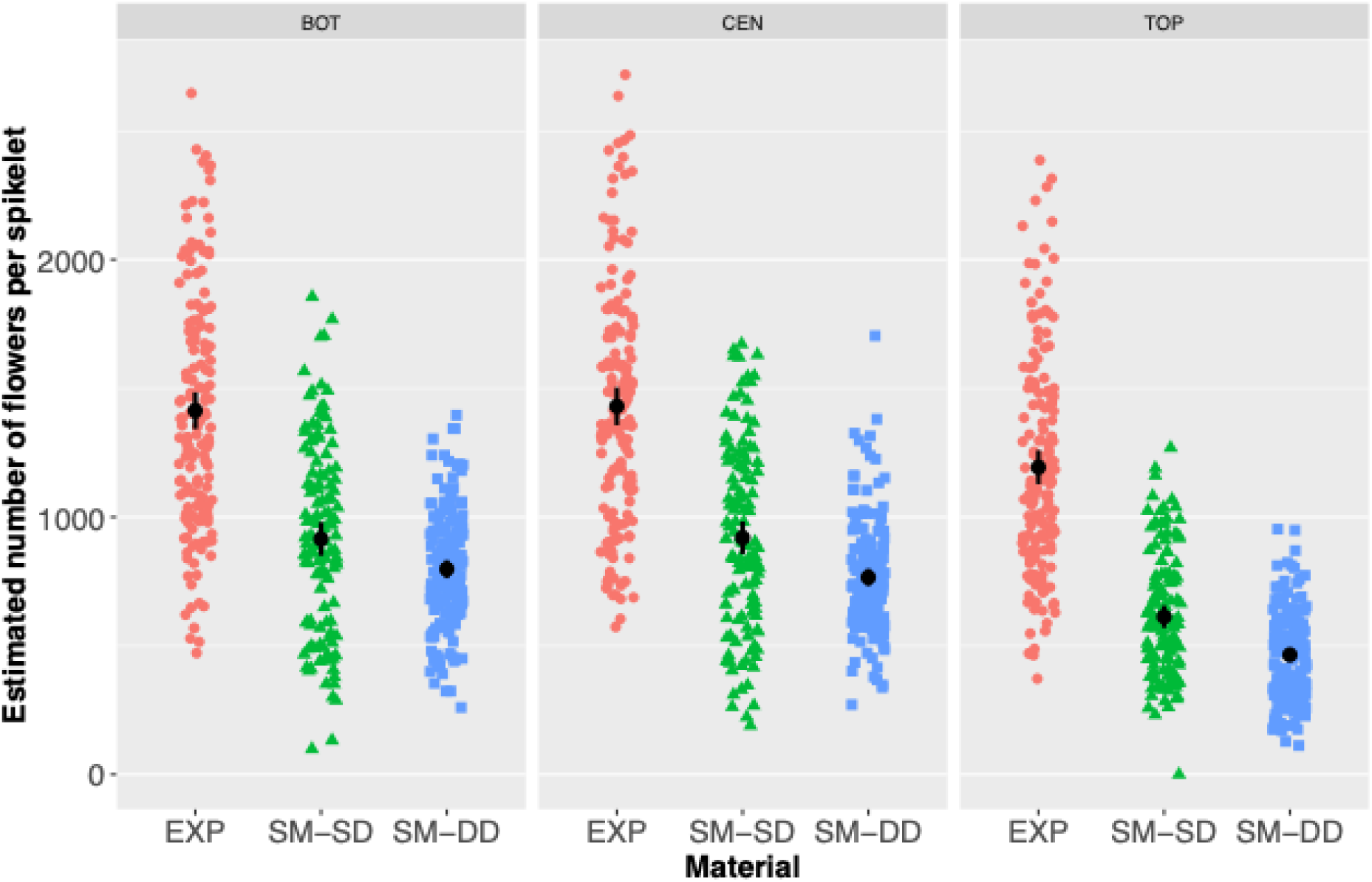
Estimated number of flowers per spikelet depending on the spikelets position on the inflorescence and the planting material.

When looking at the number of flowers per useful cm (Figure 11), a significant interaction between the planting material and the position of the spikelets was found (*p =* 7.625 × 10^−5^), indicating that the variations in the number of flowers per cm across materials were not uniform and depended critically on the specific region of the inflorescence being observed. Post-hoc pairwise comparisons revealed that the material performance varied distinctly by position (Figure 12). At both the bottom and the center positions, the EXP material exhibited significantly more flowers per cm compared to both SM materials (*p* < 0.0229), while no significant difference was observed between SM-SD and SM-DD material (*p* > 0.1667). However, a major shift occurred at the top position, with SM-SD material having a substantial increase, successfully catching up to the point of being statistically indistinguishable from EXP (*p* = 0.4232) and also SM-DD (*p*= 0.1667) materials. In contrast, SM-DD material remained significantly lower than EXP material at the top position (*p* = 0.0066).

**Figure 11.**
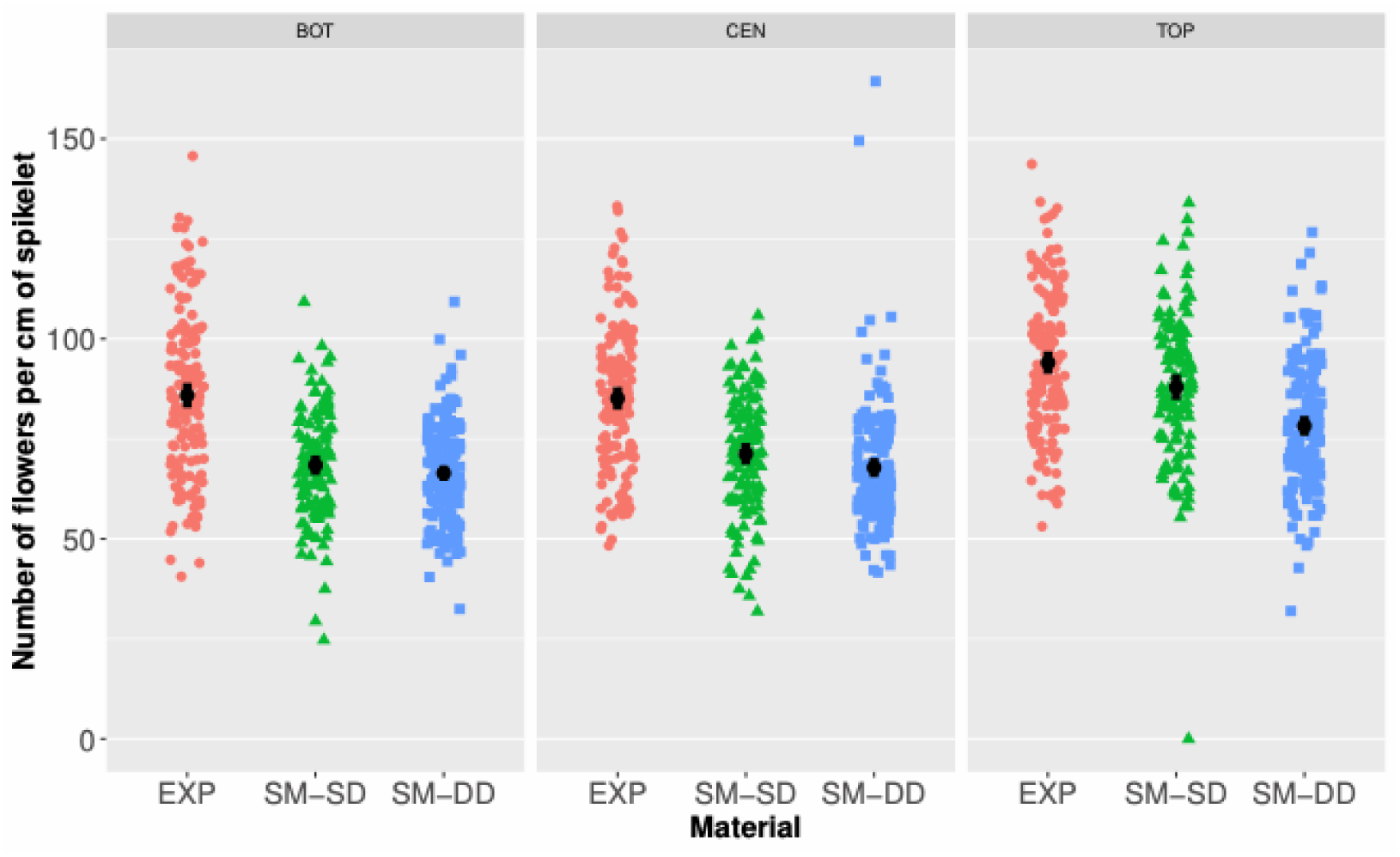
Estimated number of flowers (per cm of spikelet) depending on the spikelets position on the inflorescence and the planting material.

**Figure 12.**
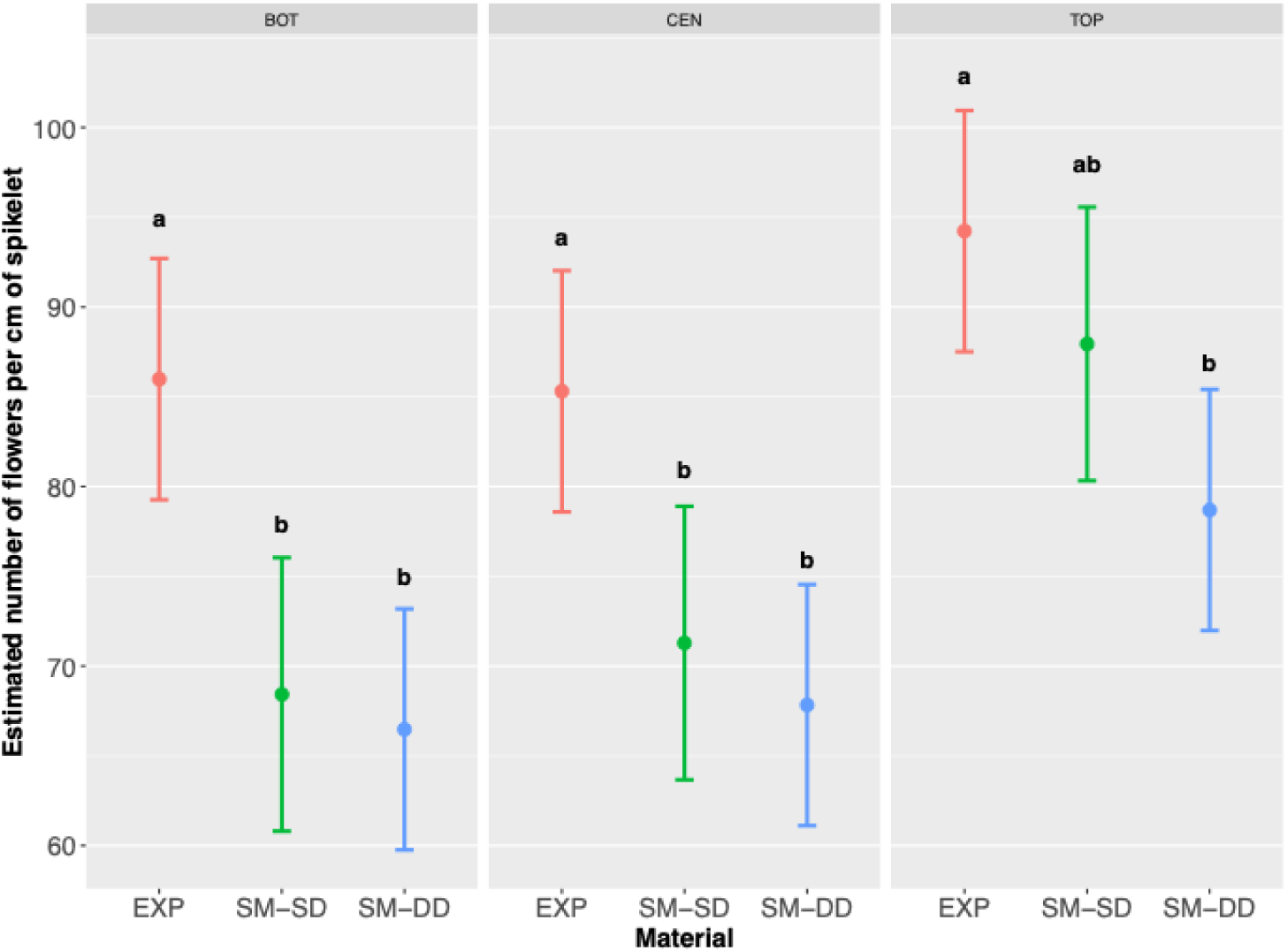
Estimated number of flowers per cm of spikelet estimated from the LMM model (with standard error). Different lowercase letters within the same inflorescence zone (BOT, CEN, TOP) indicate statistically significant differences between plant materials (Tukey’s HSD test, p < 0.05)

Finally, when taking the useful length and the number of flowers per spikelet into account we estimated the total number of flowers of the whole inflorescence for all 3 planting materials, and show that no significant difference was observed among them (Figure 13, *p* > 0.4088).

**Figure 13.**
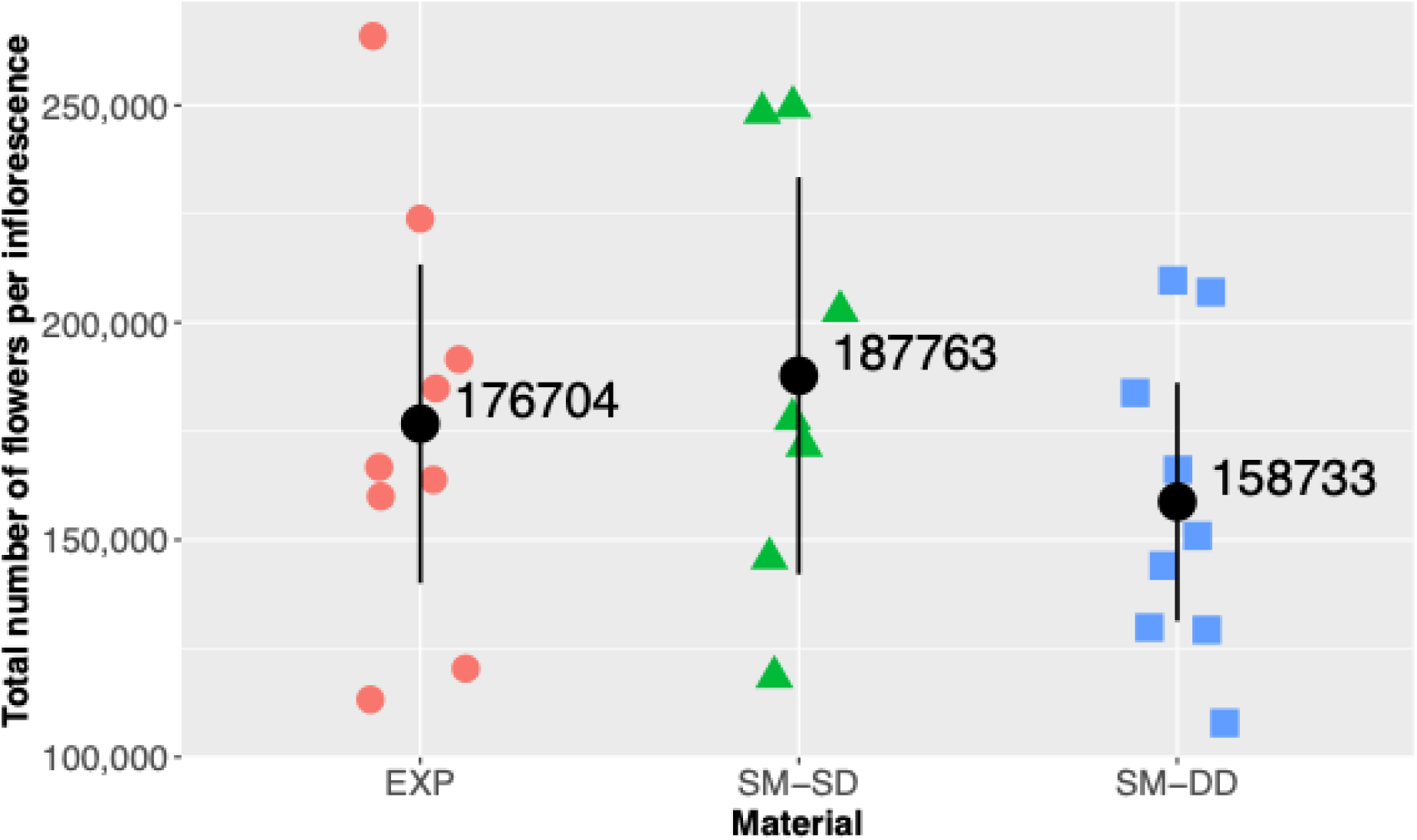
Estimates of the average number of potential flowers per inflorescence (and standard error).

### Number of flowers occupied by *E. kamerunicus* immature stages

Number of flowers per cm occupied by *EK* (Figure 14) was not significantly affected by planting material (*p* = 0.1564) and by spikelets position on the inflorescence (*p* = 0.5189). No significant interaction was found here (*p* = 0.6032). Post hoc comparisons showed that *EK*. occupied more flowers in EXP than in SM-DD materials (*p* = 0.0226), but there were not statistical differences between EXP and SM-SD materials (*p* = 0.376), and between SM materials (*p* = 0.4957).

**Figure 14.**
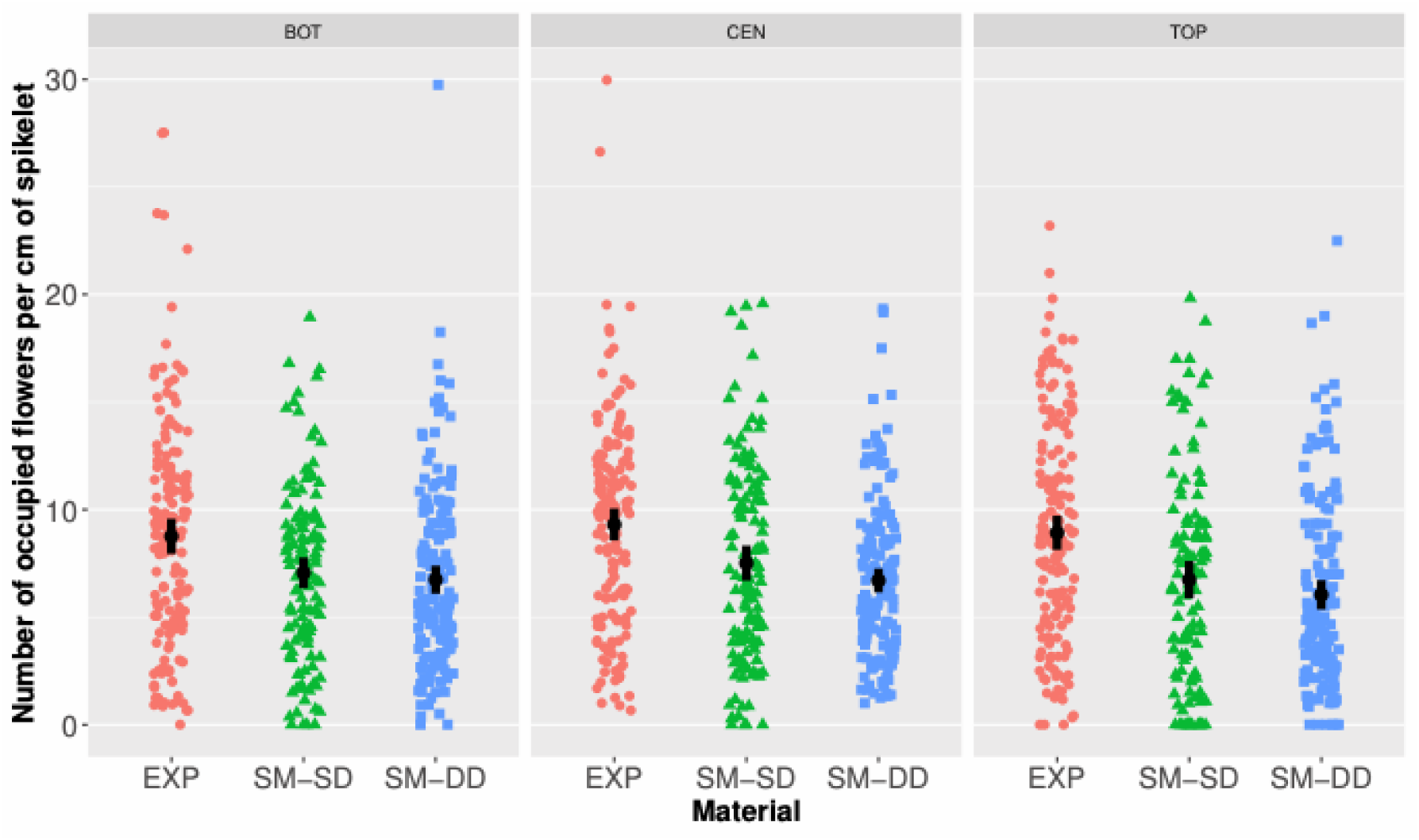
Estimated number of occupied flowers per cm of spikelet for each planting material and each position.

A post-hoc pairwise comparison (Figure 15) revealed that the planting material and the position within the inflorescence had no significant impact on the number of occupied flowers per cm of spikelet (p > 0.277). Finally, when looking at the total number of occupied flowers by immature stages of *E. kamerunicus* per inflorescence (eggs, larvae, prepupae and pupae) at the inflorescence level (Figure 16), no significant difference was observed among all 3 planting materials (*p* > 0.4884).

**Figure 15.**
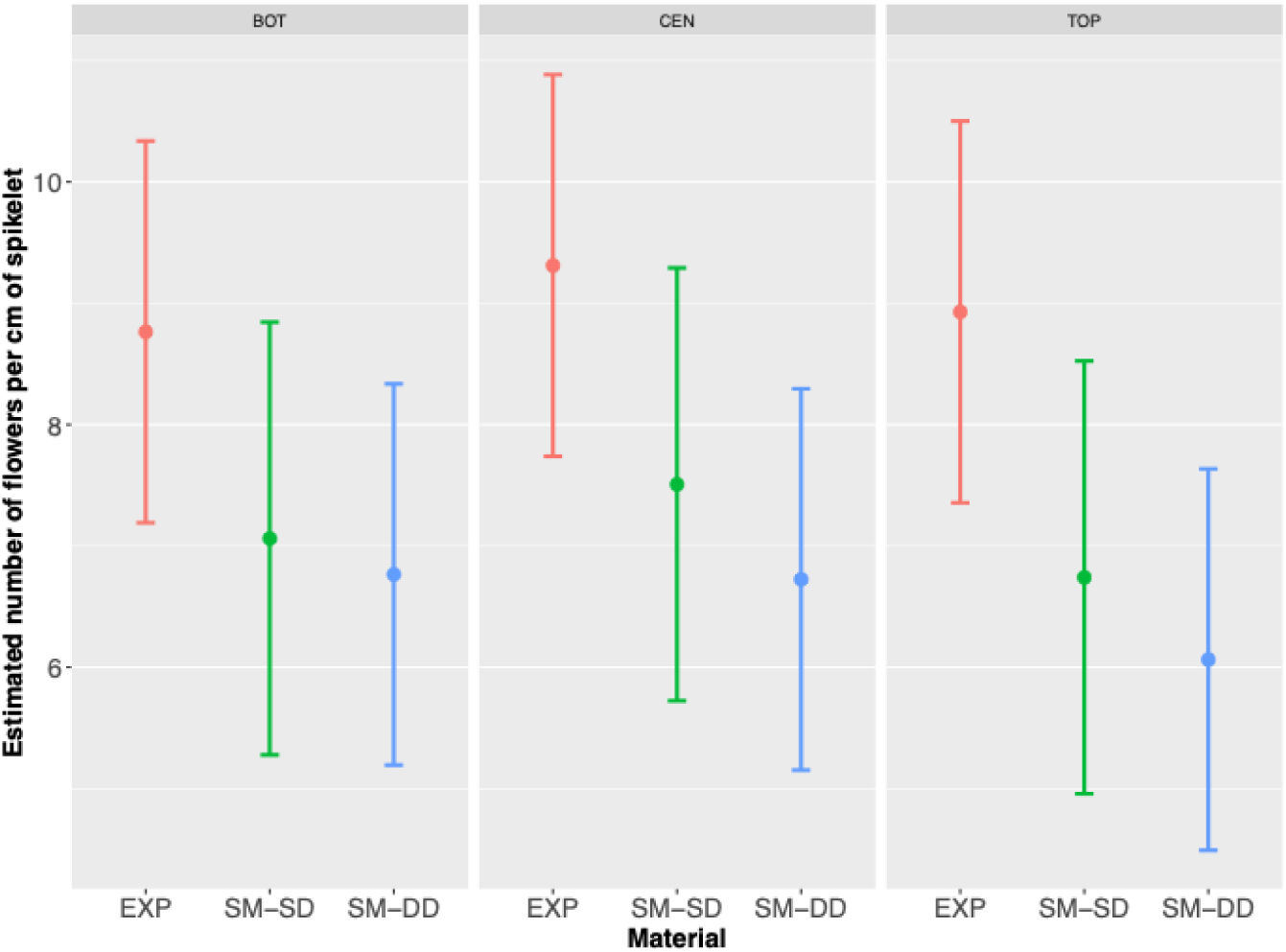
Estimated number of occupied flowers per cm of spikelet estimated from the LMM model (with standard error).

**Figure 16.**
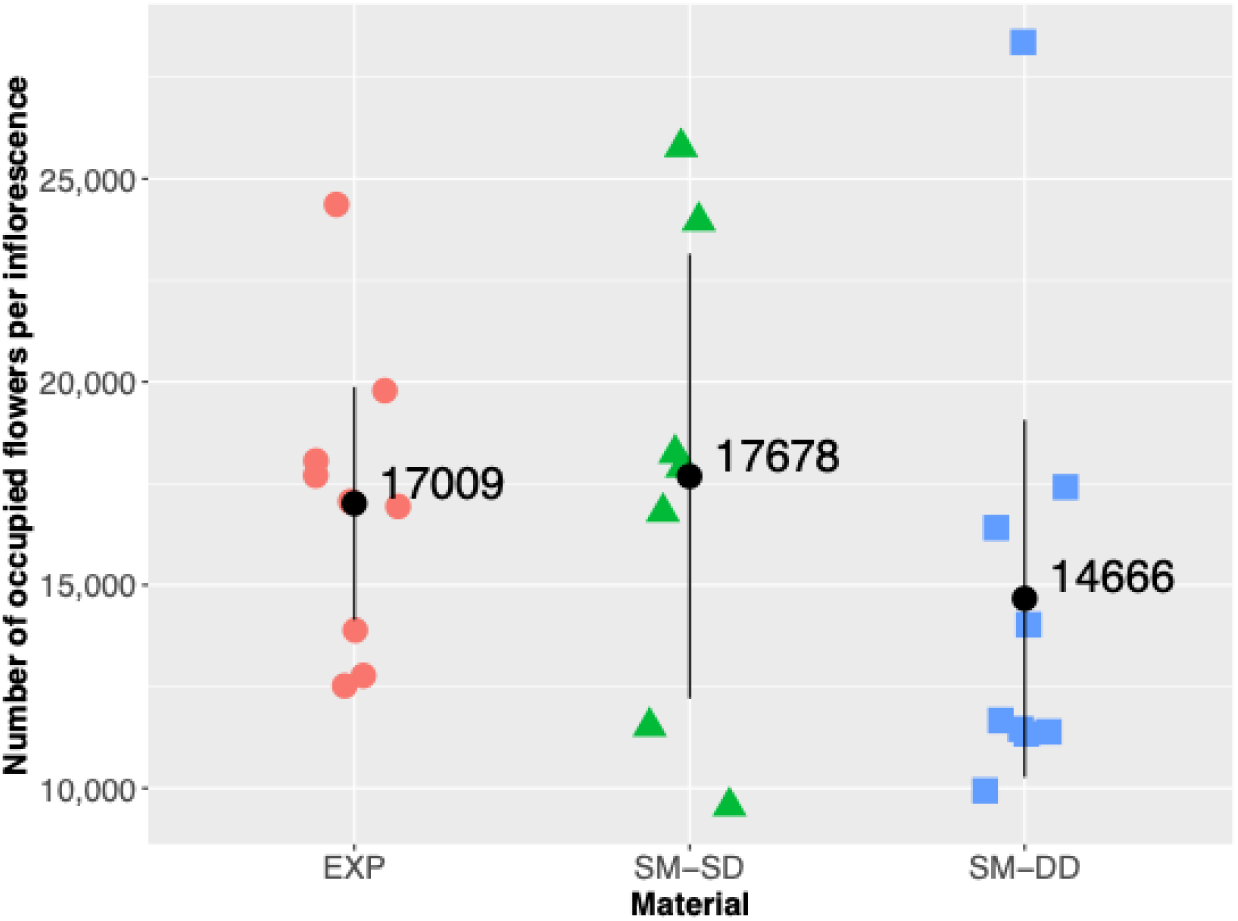
Estimates of the average number of flowers occupied by immature stages of Elaeidobius

Using data presented in Figure 13 and Figure 16, the occupancy ratio of *E. kamerunicus* was estimated to 10% overall, with 10.2 % for the EXP, 9.6 % for the SM-SD, and 9.2 % for the SM-DD material.

Adjusted means (±SE) for all variables are presented in Table 2, below, for the 3 different planting materials.

**Table 2.**
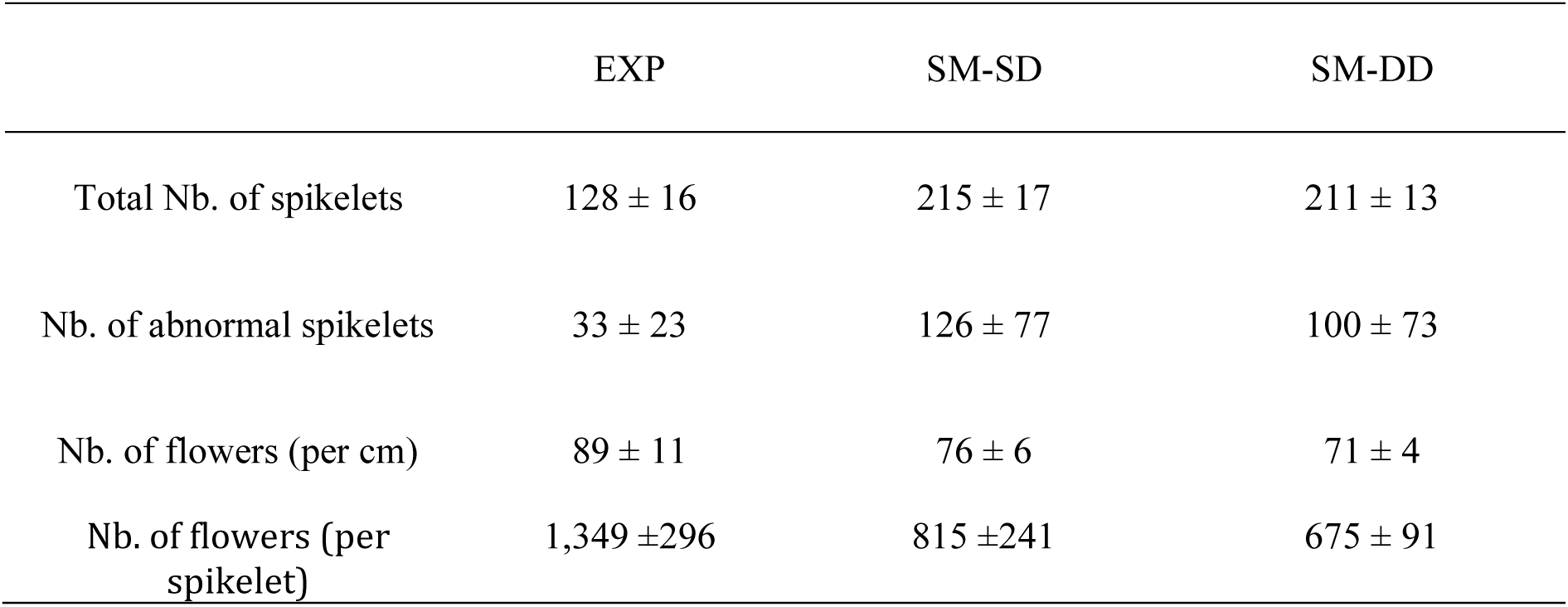
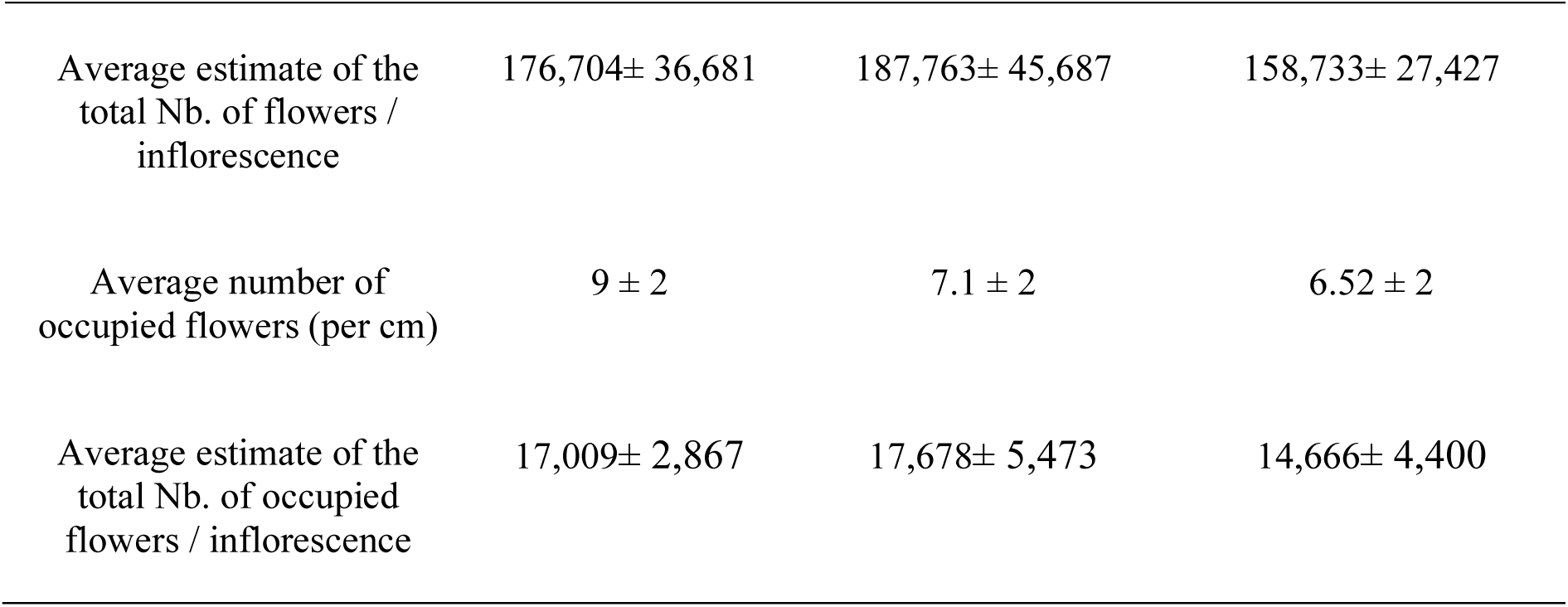
Summary of the results for the comparative study of three planting materials.

### *E. kamerunicus* immatures stages at different stages of post-anthesis male inflorescence

According to Figure 17, eggs were present during stages M_5_ and M_6_ but were almost absent at stages M_7_ and M_8_. Larvae mainly present in M_5_ and M_6_ melt into prepupae in M_7_, M_8_ and M_9_. From M_8_ onwards, prepupae melt into pupae. Comparison tests showed that the differences were highly significant (*p* = 5 × 10^−5^). This was confirmed by a Pearson standardized residual analysis, which showed that the stages development was faster in the EXP and SM-SD materials than in the SM-DD material.

**Figure 17.**
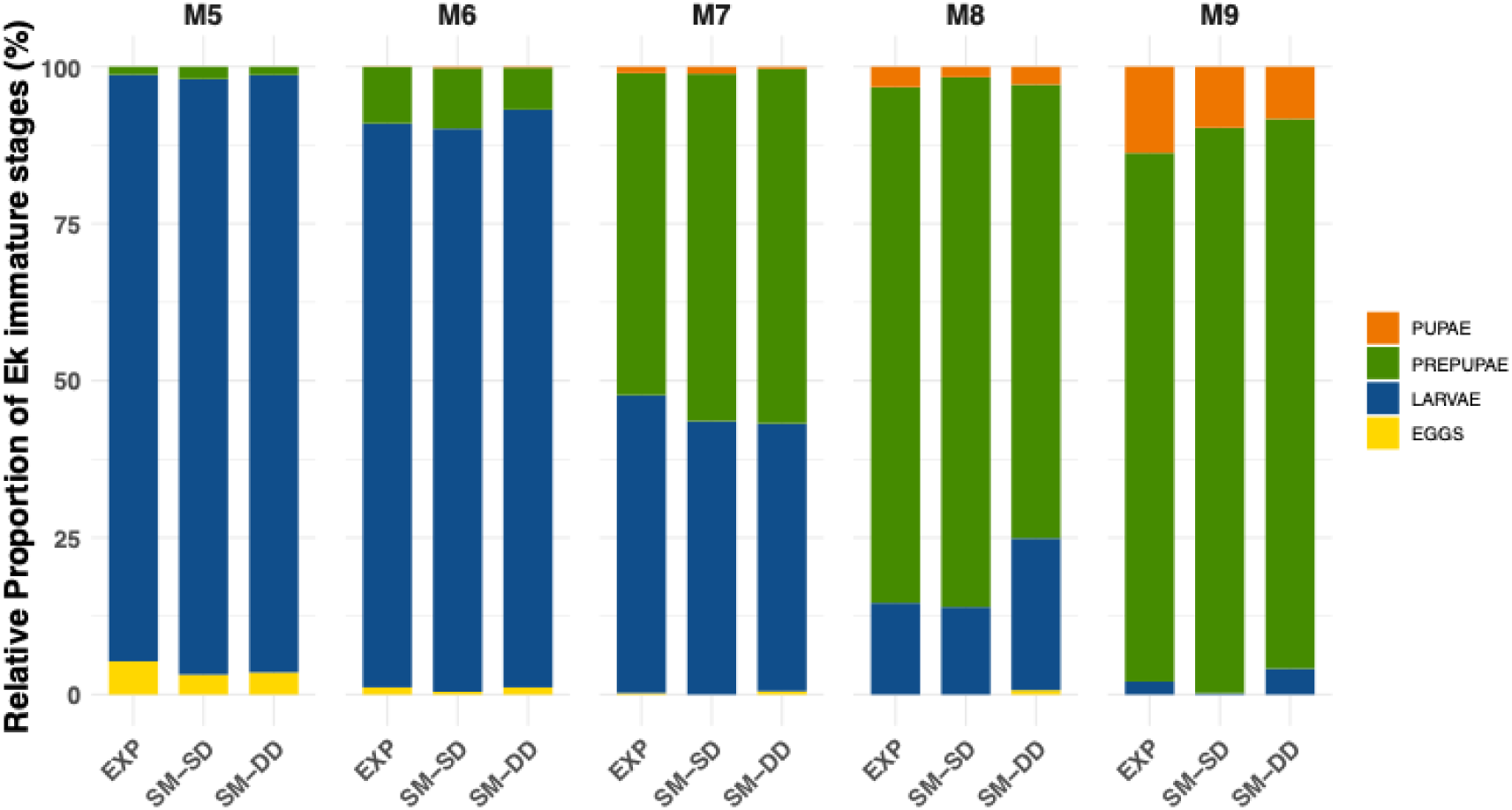
Development of E. kamerunicus immature stages in the post-anthesis male inflorescence

## Discussion

While SM planting material produced more spikelets (average of 213 ± 15 per inflorescence) than EXP planting material (128 ± 16 per inflorescence), the number of flowers per inflorescence available to host *E. kamerunicus* immature stages was similar between EXP and SM materials.

Independently of the planting material, EXP or SM, the average number of spikelets was consistent with the estimate given by (Syed, 1979; Janick and Paull, 2008; Jacquemard, 2011) between 100 and 300 spikelets per inflorescence. (Swaray et al., 2021) reported 138.41 as the mean number of spikelets per male inflorescence. According to (Hartley, 1988), the number of spikelets per inflorescence varied greatly from palm to palm. (Corley and Gray, 1976) showed that average spikelet number increased with palm age, from an average of about 60 in 3-year-old palms to 150 by 10 years. Inflorescences examined from 1,476 palms in Zaire with 125-165 spikelets were most frequent whereas the range was 85-285 spikelets (Hartley, 1988). A range from 100 to 283 has been recorded on 37-year-old palms in Congo (Corley and Tinker, 2016). The older oil palms were, the greater was the number of spikelets as also indicated by (Rahardjo et al., 2018; Tandon et al., 2001). The male inflorescence became larger in size as the oil palm ages, regarding both the length of individual spikelets and the number of spikelets per inflorescence (Corley and Tinker, 2016).

Nevertheless, the average number of spikelets was still higher in SM (213 in average) that in EXP palms (128 on average) which might compensate for its large number of abnormal spikelets (100 and 126 in SM in double density and single density respectively against 33 in EXP palms). Abnormal spikelets could have a negative effect on the useful area of flowers available for egg-laying for the development of *E. kamerunicus*. The estimates of useful area showed significant parity between EXP and SM. Indeed, the higher total number of spikelets in SM compensated the negative effect due to abnormal spikelets. It was important to note that abnormal spikelets characterized by part of the spikelet without any flowers was described for the first time here from SM planting material.

According to (Kurniawan, 2010; Rahmadani and Iqba, 2015), the number of spikelets per male inflorescence at anthesis affected the number of *E. kamerunicus* populations. (Kouakou et al., 2018; Nurul Fatihah et al., 2019) reported the number of male inflorescences and spikelets were positively correlated with the adults *E. kamerunicus* population per spikelet and were the main attributes for attracting a higher number of *E. kamerunicus* (Yousefi et al., 2020). Similar results were obtained by (Syarifah and Idris, 2016). For (Swaray et al., 2021), there was a moderate positive significant correlation.

The EXP spikelets contained an average of 1,349 male flowers whereas the SM spikelets contained 815 and 675 male flowers for single and double density respectively. Our findings were consistent with those reported by (Janick and Paull, 2008; Mohamad et al., 2021), who reported that each of the spikelets usually holded 400-1,500 sessile male flowers. According to (Tan and Rao, 1979), spikelets of 8-year-old tenera palms in Malaysia were found to have an average of 785 flowers. With about 150 spikelets, there would thus be well over 100,000 flowers per inflorescence (Corley and Tinker, 2016). The spikelet of a medium size inflorescence beared about 750 flowers, so that a mature inflorescence with 200 spikelets will bear 150,000 male flowers (Turner, 1974). This observation was more consistent with our estimate of 158,733 per inflorescence for SM planted in double density.

In the present study, we found 76 and 71 flowers per cm (SM-SD and SM-DD respectively) and 89 flowers per cm for EXP. Our results showed that these planting materials produced much more flowers than the material considered in (Ponnamma et al., 2006) for which the number of flowers per cm ranged from 40.21 to 55.14.

Furthermore, regardless of the material, there was a significative difference in terms of useful area (in cm) between the top and the rest of the inflorescence. Spikelets located at the top had more flowers per cm. The explanation might lie in the average diameter of the spikelets, which was greater at the top of the inflorescences. These average diameters ranged from 1.1 cm to 1.5 cm and were similar to those reported by (Corley and Tinker, 2016) ranging from 0.8 cm to 1.5 cm. Even though the number of flowers (per cm) was greater at the top of the inflorescence, the *E. kamerunicus* occupancy rate was lower compared to the bottom and the center of the inflorescence. (Mohamad et al., 2021) reported that the length of spikelets sampled from the top of the inflorescences was generally shorter compared to samples collected from the other parts of the inflorescences. Shorter spikelets would have less flowers and might influence the amount of food and breeding sources available as also noted by (Ming et al., 2022). The same author stated in other months of samplings, no significant differences were found between the length of spikelets from different portions of male inflorescence.

According to (Turner and Gillbanks, 1974), flowering began gradually from the base of the inflorescence rachis to its apex. When flowers open, the fragrance of estragole released from the male flowers (Syed et al., 1982; Anggraeni et al., 2013) attracted *E. kamerunicus* (Chiu et al., 1986). As these attractive molecules were released throughout anthesis, this would imply that at the beginning of anthesis (stage M_0_ to stage M_3_), *E. kamerunicus* would be preferentially attracted to the base and center of the inflorescence without being attracted to the top. Consequently, it was likely that *E. kamerunicus* had the choice of laying their eggs at the base or the center of the inflorescence rather than at the top for a longer period of time. This could explain why the occupancy rate of the lodges (per cm) was lower at the top of inflorescences despite a higher number of available flowers. Furthermore, the high insect concentration within few inflorescences might have a negative impact on the species, since it might lead to an increase in mortality rate and a greater risk of disease transmission (Syed, 1978).

The oviposition period ranged from 7-12 days (Dhileepan, 1994). According to our results, the evolution of immatures stages of *E. kamerunicus* inside spikelets showed the development from M_5_ to M_10_ of the post-anthesis stage. It was similar for both planting materials, EXP and SM. The mean number of immature stages was increasing for 4-5 days and the decrease due to mould (Prasetyo et al., 2013). According to (Syed, 1982), the minimum larval development into pupae was 5 days and the pupae to imago at least 2 days. (Kurniawan, 2010) found a minimum of 6 days and 3 days in West Java and Central Kalimantan respectively.

We found on average 10% of the flowers occupied by *E. kamerunicus* immatures stages in both planting material. Our findings meaned that the estimate number of immature *E. kamerunicus* counted during this study was lower than the potential occupancy of the inflorescence. On average, we found 180,000 flowers per male inflorescence and 18,000 occupied by *EK*. (Syed, 1982) mentioned that one individual might visit up to six flowers to complete its larval development. This would mean theoretically a maximum potential of 30,000 flowers occupied by *EK* immature stages (16.67 %). According to (Prasetyo and Susanto, 2012), if an *E. kamerunicus* larva consumed 3 to 5 flowers and if a male inflorescence contained 100,000 flowers, 20,000 *E. kamerunicus* larvae could develop (20%). (Syed and Saleh, 1988) stated the entire food source (male flowers) was not fully utilised. The percentage of occupied flower indicated that male inflorescence in anthesis as host was not at optimum levels due to a low weevil population (Prasetyo et al., 2014). A range of 3.71 to 25.88 larva per cm was reported by (Ponnamma et al., 2006). This result of flowers occupancy was also very useful to parameterize mathematical models based on non-adult carrying capacity per male inflorescence. It confirmed the carrying capacity value used in (Dumont et al., 2018) and will be useful for further modelling analysis and simulations.

(Syed, 1982) stated *Elaeidobius* spp. did not oviposit inside flowers of any other plants except oil palm. Host preference was associated with higher oviposition rates, increased female longevity, shorter development time and higher survival in all life stages (Lenteren and Noldus, 1990). Biotic and abiotic factors could influence population pollinator weevils (Lubis et al., 2023; Zulkefli et al., 2020). During the present study, rat attacks on male inflorescence were observed in the field. (Ponnamma et al., 2006) and (Saravanan et al., 2024) stated rats might affect *E. kamerunicus* population, searching of eggs, larvae, pupae and imago, preyed upon them and causing high mortality rate. (Lubis et al., 2023) reported an average of 140 weevil larvae per spikelet in absence of rat attack to male inflorescence.

According to (Nurul Fatihah et al., 2019), the rainy season was suitable to *E. kamerunicus* development. (Dhileepan, 1994; Hutauruk and Ps, 1984; Syed and Saleh, 1988) and (Wahid et al., 1987) reported similar observations. Not on *E. kamerunicus* development but weevil population and weevil pollination activity, (Sugih et al., 1996) reported wetter weather possibly having negative effects. The present study occurred during a relatively dry season. The rainfall data recorded did not exceed 400 mm from April to June 2019, so the impact of heavy rainfall on the population was not tested. Moreover, the weevil population size in the male inflorescence differed significantly between months (Dhileepan, 1994).

Application of various insecticides directly on male inflorescence at different anthesis stages affected breeding of *E. kamerunicus* by decreasing the number of new emerged weevil from each spikelet of male inflorescence (Ismail et al., 2020; Ming and Bong, 2017; Prasetyo and Susanto, 2019; Setyawan et al., 2020). Other interactions such as predation and parasitism are not yet well understood yet (Li et al., 2019).

Our results gave an accurate estimate of the immature weevil population on post-anthesis male inflorescences before *E. kamerunicus* imago emerged. It considered the time of sampling, different stages post-anthesis and stratified sampling of spikelets on the male inflorescence.

From many previous studies, estimation of the weevil population was obtained by counting the number of weevil present on spikelets on different part of the inflorescence in anthesis (Basri et al., 1987; Dhileepan, 1994; Nurul Fatihah et al., 2019; Swaray et al., 2021; Lubis et al., 2023).

## Conclusion

The goal of developing “super-male” palms was to compensate for the hyperfeminization of the EXP material by producing a greater number of male inflorescences. However, it was necessary to ensure that the male inflorescences were equivalent—from the pollinator’s perspective—to the male inflorescences of the EXP material. Our findings indicated the equivalent potential of male inflorescence of SM and EXP to host *E. kamerunicus* immature stage development, despite noticeable morphological differences.

These first estimates of the immature stages occupying male inflorescences were interesting in light of future estimation of *E. kamerunicus* progeny measured from the spikelets of EXP compared to those from SM inflorescences. Thus, the relationship between the male inflorescence and the weevil population needs to be further investigated to determine the potential of male inflorescence to support the development of *E. kamerunicus* imagos responsible of the pollination. In the meantime, our results raised a question which would be to understand why the estimated number of *E. kamerunicus* immature stages per inflorescence does not reach the full potential of the inflorescence and how it may, eventually, fluctuate along the year.

## Data

All data will be available at: https://doi.org/10.18167/DVN1/MCBBTL

## Acknowledgments

The authors would like to acknowledge funding from Palmelit SAS, Cirad and PT Socfindo Indonesia. Special thanks to Mr N. Turnbull who provided assistance in the ELAPALM project. Finally, we particularly appreciate the support of PT Socfindo Indonesia executive director, Mr H. O. Williams for fieldwork facilities and research permissions.

## Author Contribution

LBO, HR, and YD conceived and designed research. CM, IS and DA supervised the observations. EF, RRM made the initial observations. EF made the preliminary analysis. YD performed the statistical analyses. YD and LBO wrote the original draft.

